# Nogo-A Beyond the RhoA/ROCK Pathway – Novel Components of Intracellular Nogo-A Signaling Cascades

**DOI:** 10.1101/2020.07.10.197368

**Authors:** Martina A. Maibach, Ester Piovesana, Julia Kaiser, Mea M. Holm, Zorica Risic, Michael Maurer, Martin E. Schwab

**Affiliations:** Brain Research Institute and Department of Health Sciences and Technology, University and Dept. of Health Sciences and Technology, ETH Zurich, 8057, Zurich, Switzerland

## Abstract

Nogo-A is a well-characterized myelin-associated membrane protein that restricts fibre growth and the regenerative capacity of the adult central nervous system after injury. To date Nogo-A post-receptor signalling pathway research focused on the RhoA/ROCK cascade, which can lead to growth cone collapse and neurite retraction. Much less is known about continued intracellular Nogo-A signalling mediating long-term neurite outgrowth inhibition resulting from transcriptional and translational changes. Here, we propose a simple but highly reproducible *in vitro* assay to study Nogo-A related signaling and neurite outgrowth inhibition in general. Furthermore, we identified ERK1/2 as downstream effector of Nogo-A, partially mediating its neurite outgrowth inhibition. We describe ERK1/2 dependent changes of translational events such as elevation of RhoA levels within the growth cone, which may potentiate the cells’ responses to Nogo-A. We also observed Nogo-A dependent upregulation of the JAK/STAT pathway inhibitors SOCS3 and KLF4 and downregulation of insulin mediated phosphorylation of AKT, indicating direct negative crosstalk between Nogo-A signalling and the growth promoting JAK/STAT and AKT/mTORC1 pathways.

## Introduction

Following injury or trauma, the central nervous system (CNS) has a low regenerative capacity, in part due to the growth limiting environment containing axonal growth restricting factors such as myelin-associated inhibitors, scars and perineuronal nets (Schwab and Strittmatter, 2014). One of the most extensively studied myelin associated neurite growth inhibitors is the membrane protein Nogo-A. Antibody mediated neutralization, knock out or pharmacological inhibition of Nogo-A or its receptors leads to enhanced sprouting of injured and spared axons and to higher levels of functional recovery in several models of CNS lesion, e.g. spinal cord injury and stroke (Schwab and Strittmatter, 2014).

Nogo-A signals via two functional domains, Nogo-A-Δ20 and Nogo-66, which bind to several domain-specific, signal transducing receptors (Kempf and Schwab, 2013). The known functional receptors for the Nogo-66 domain are Nogo receptor 1 (NgR1), paired immunoglobin-like receptor B (PirB) and low density lipoprotein receptor (LRP) (Kempf and Schwab, 2013). Only recently sphingosine-1-phosphate receptor 2 (S1PR2), tetraspanin 3 (TSPAN3) and syndecan 3/4 (SDC3/4) have been identified as receptors for the Nogo-A-Δ20 domain (Kempf et al., 2017; Thiede-Stan et al., 2015; Kempf et al., 2014; Kempf and Schwab, 2013). Interestingly, both Nogo-A-Δ20 and Nogo-66 signaling induce the activation of the small GTPase Ras homologue A (RhoA) and its downstream effector Rho-associated, coiled-coil containing protein kinase (ROCK), which mediates cytoskeletal rearrangements leading to growth cone collapse and retraction. Apart from this immediate downstream signaling, Nogo-A can also induce transcriptional changes (Craveiro et al., 2008; Bareyre et al., 2002; Buffo et al., 2000), suggesting continued signaling to the cell body and nucleus. The molecular basis of this retrograde signaling is still largely unknown.

In the context of CNS trauma, the protein Kinase B (AKT)/mammalian target of rapamycin complex 1 (mTORC1) and janus kinase (JAK)/signal transducer and activator of transcription protein (STAT) cascades have been associated with promoting axonal regeneration (Lu et al., 2014). Deletion or pharmaceutical inhibition of phosphatase and tensin homolog (PTEN), an upstream inhibitor of the AKT/mTORC1 pathway (Saxton and Sabatini, 2017), results in enhanced axonal regeneration following optic nerve crush and spinal cord injury (Danilov and Steward, 2015; Du et al., 2015; Lewandowski and Steward, 2014; Ohtake et al., 2014; Zukor et al., 2013; Park et al., 2008). Similarly, activation of the JAK/STAT signaling cascade via deletion of its inhibitors suppressor of cytokine signaling 3 (SOCS3) or kruppel like factor 4 (KLF4) increased retinal ganglion cell regeneration after optic nerve crush (Moore et al., 2009; Smith et al., 2009). The activation of these pathways is regulated by a highly interconnected signaling network integrating both cell external and internal cues (Mazel, 2017; Saxton and Sabatini, 2017; Rawlings, 2004). So far, it has not been investigated whether inhibitory factors such as Nogo-A impact on or influence these pathways directly, in part due to the lack of a suitable system to screen for potential modifications.

In this study, we propose a simple but highly reproducible *in vitro* assay to study Nogo-A related signaling and neurite outgrowth inhibition in general. We identify extracellular signal regulated kinase (ERK1/2) as an effector propagating Nogo-A signaling. We describe ERK1/2 downstream events such as elevation of RhoA levels, which may enhance Nogo-A mediated growth cone collapse and retraction. We also observe a signaling crosstalk between Nogo-A and the axonal growth promoting AKT/mTORC1 and JAK/STAT pathways at multiple levels. Taken together, these results suggest that Nogo-A signaling not only leads to growth cone collapse and retraction via the RhoA/ROCK axis, but also results in inhibitory crosstalk with growth-promoting cascades.

## Methods

### Experimental Animals

All animal experiments were performed with the approval of and in strict accordance with the guidelines of the Zurich Cantonal Veterinary Office. All efforts were made to minimize animal suffering and to reduce the number of animals required.

### Spinal cord extracts

Rats were decapitated, and the spinal cords were dissected, followed by immediate homogenization in ice-cold extraction buffer (3.7% CHAPS and 5 mM EDTA in PBS) containing 1x HALT ™ protease and phosphatase inhibitor (Thermo FisherScientific). The homogenates were incubated for 30 min on ice, and centrifuged 4 times for 15 min at 13’000 g at 4°C. The supernatants were aliquoted, snap-frozen in liquid nitrogen, and stored at -80 °C until use.

### Antibodies and reagents

For neurite outgrowth assays the following molecules were used at the indicated concentrations: 10 µg/ml mouse anti-Nogo-A antibody 11C7 (produced in house; Oertle et al. (2003)), 10 µg/ml mouse anti IgG1 M-BSV-1 (APS, FG12/B5), 1 nM JTE-013 (Tocris, 2392), 0.5 U Heparinase III (Sigma, H8891), 2 nM NEP1-40 (Sigma, N7161), 1 nM SCH772984 (Selleckchem, S7101), 10 ng/ml IL-6 (R&D, 406-ML-005). For western blotting, the primary antibodies are summarized in table Table 1.1. Secondary HRP-coupled antibodies were all purchased from Thermo Fisher Scientific and used at a concentration of 0.05-0.1 μg/ml. For immunocytochemistry, the following primary antibodies were used at the indicated concentrations: 1:10’000 mouse anti-β3-Tubulin (Promega, G712A), rabbit anti-p-ERK T202/Y204 (CST, 9101), mouse anti-RhoA (Santa Cruz, SC-418). As secondary antibodies, 1μg/ml anti-mouse or anti-rabbit Cy3-coupled antibodies (Invitrogen, A10521, A10520) were used. 1:100 Phalloidin Alexa Fluor488 (Invitrogen, A12379) was used to stain the f-actin cytoskeleton, and 50 nM DAPI (Invitrogen, D3571) was used as a nuclear counterstain.

**Table 1.1.** Antibodies used for Western Blotting.

| Target | Host Species | Dilution | Company | Catalogue Number |
| --- | --- | --- | --- | --- |
| p-AKT S473 | rabbit | 1:1000 | Cell Signaling Technology | 9271 |
| p-AKT T308 | rabbit | 1:500 | Cell Signaling Technology | 9270 |
| AKT | rabbit | 1:1000 | Cell Signaling Technology | 9272 |
| p-4EBP1 S65 | rabbit | 1:500 | Cell Signaling Technology | 9451 |
| p-4EBP1 T37/46 | rabbit | 1:500 | Cell Signaling Technology | 2855 |
| 4EBP1 | rabbit | 1:500 | Cell Signaling Technology | 9644 |
| p-S6K S389 | mouse | 1:1000 | Cell Signaling Technology | 9206 |
| S6K | rabbit | 1:1000 | Cell Signaling Technology | 2708 |
| p-S6P S235/236 | rabbit | 1:1000 | Cell Signaling Technology | 4858 |
| S6P | mouse | 1:1000 | Cell Signaling Technology | 2317 |
| p-ERK1/2 T202/Y204 | rabbit | 1:2000 | Cell Signaling Technology | 9101 |
| ERK1/2 | rabbit | 1:1000 | Cell Signaling Technology | 4695 |
| p-RSK S380 | rabbit | 1:1000 | Cell Signaling Technology | 11989 |
| RSK | rabbit | 1:1000 | Cell Signaling Technology | 9355 |
| RhoA | rabbit | 1:1000 | Cell Signaling Technology | 2117 |
| p-CREB S133 | rabbit | 1:1000 | Cell Signaling Technology | 9198 |
| CREB | rabbit | 1:1000 | Cell Signaling Technology | 9197 |
| RhoA | rabbit | 1:1000 | Cell Signaling Technology | 2117 |
| p-c-Myc S62 | rabbit | 1:1000 | Cell Signaling Technology | 13748 |
| c-Myc | rabbit | 1:10'000 | Abcam | ab32072 |
| c-Jun | rabbit | 1:500 | Cell Signaling Technology | 9165 |
| p-STAT3 S727 | rabbit | 1:500 | Cell Signaling Technology | 94994 |
| STAT3 | mouse | 1:1000 | Cell Signaling Technology | 9139 |
| SOCS1 | rabbit | 1:1000 | Cell Signaling Technology | 3950 |
| SOCS3 | rabbit | 1:1000 | Cell Signaling Technology | 2932 |
| KLF4 | rabbit | 1:1000 | Cell Signaling Technology | 12173 |
| GAPDH | mouse | 1:20'000 | Abcam | ab8245 |

### Cell Culture

N1E-115 mouse neuroblastoma cells were obtained from ATCC and maintained at 37 °C in a humidified atmosphere with 5% CO_2_ in DMEM (Sigma) supplemented with 10% FBS (Sigma), 2% L-Glutamine and 1% P/S. Neuron like differentiation was induced by switching the medium to Neurobasal (Gibco) supplemented with 2% L-Glutamine (Sigma) and 1% P/S (Sigma).

### Neurite outgrowth assay

Cells were plated in differentiation medium at a density of 10’000 cells/cm^2^. After 24 h the cells were treated with spinal cord extract and molecular factors and left to grow for another 24 h. The assays were stopped by the addition of 4% PFA at RT for 15 min. The cells were then counter-stained with Coomassie solution (0.25% Coomassie Brilliant Blue R250 (Sigma), 50% MeOH, 10% AcOH) for five minutes, followed by two consecutive washes with PBS and stored at 4°C. Four images at predefined locations in the wells were acquired on a ScanR HCS microscope (Olympus) equipped with an UPLSAPO dry 10x/0.4 objective and Hamamatsu ORCA-FLASH 4.0 V2 camera using the ScanR analysis software. Mean neurite outgrowth was quantified in ImageJ by applying a grid to the pictures and counting intersections of neurites with the grid lines and total cell bodies and calculating the ratio thereof (Ronn et al., 2000). Each experiment was conducted in three technical replicates and all quantifications show five independent experiments.

### Cell Lysis

Cells were washed twice in PBS on ice and lysed in RIPA buffer (150 mM NaCl, 1% NP-40, 1% Sodium deoxycholate, 0.1% SDS, 50 mM Tris pH8) containing 2x HALT™ phosphatase inhibitor cocktail and 5 mM EDTA. The lysates were incubated on ice for 30 min and centrifuged at 13’000 g for 15 min at 4 °C. The supernatants were collected and stored at –80°C.

### Immunoblotting

The samples were prepared in Laemmli buffer (Biorad) supplemented with 10% βMEtOH and denatured at 90°C for 3 min. The samples were separated with pre-cast 4-15% Mini PROTEAN R TXG TM gels (Biorad) at 250 V in Tris-Glycine running buffer (25 mM Tris, 192 mM Glycine, 0.1% SDS, pH 8.3). Proteins were transferred onto a 0.45 µm PVDF membrane in Tris-Glycine transfer buffer (25 mM Tris, 19 2mM Glycine, 20% MeOH) for 90 min with a constant current of 300 mA. Subsequently, membranes were blocked for 1 h with 5% BSA (Sigma) in TBS-T (10 mM Tris, 150 mM NaCl, 0.01% Tween-20, pH 7.5) and probed with primary antibodies overnight at 4°C. The membranes were washed 3 times in TBS-T, probed with secondary HRP-coupled antibodies for 1 h at RT and washed again 3 times in TBS-T. Detection was performed using SuperSignal™ West PICO (Thermo Scientific) or WesternBright™ Sirius (Advansta) chemiluminescent substrates and images were acquired on the Gel Doc™ imager (Biorad). Densitometry analysis was performed with ImageJ software (NIH, Bethesda, MD, USA) and values were normalized to the housekeeping gene GAPDH or total protein and the mean value of the corresponding control group.

### Dot Blot

Serial dilutions of rat Nogo-A-Δ20 (produced in-house, Oertle et al. (2003)) and spinal cord extract were transferred onto a 0.45 µm PVDF membrane. Subsequently, membranes were blocked for 30 min with 5% milk powder (Migros) in TBS-T (10 mM Tris, 150 mM NaCl, 0.01% Tween-20, pH 7.5) and probed 1 h with 11C7 1:40’000 at RT. The membranes were washed three times in TBS-T, probed with secondary HRP-coupled antibody for 1 h at RT and washed again 3 times in TBS-T. Detection was performed using SuperSignal™ West PICO (Thermo Scientific) or WesternBright™ Sirius (Advansta) chemiluminescent substrates and images were acquired on the Gel Doc™ imager (Biorad). Densitometry analysis was performed with ImageJ software (NIH, Bethesda, MD, USA).

### qRT PCR

Total RNA from cultured cells was isolated with the RNeasy Micro kit (Qiagen) according to manufactures instructions and reverse transcribed using TaqMan Reverse Transcription reagents (Applied Biosystems). cDNA was amplified using the Light Cycler 480 thermocycler (Roche) with the polymerase ready mix (SYBR Green I Master, Roche). Relative quantification was performed using the comparative CT method (Schmittgen and Livak, 2008) and cDNA levels were normalized to the reference gene TBP. Each reaction was done in triplicates and quantification shows data from three independent experiments. The primers were validated using melting curve analysis of PCR products followed by gel electrophoresis to verify the amplicon size. The primers used in this study are outlined in table 1.2.

**Table 1.2.**
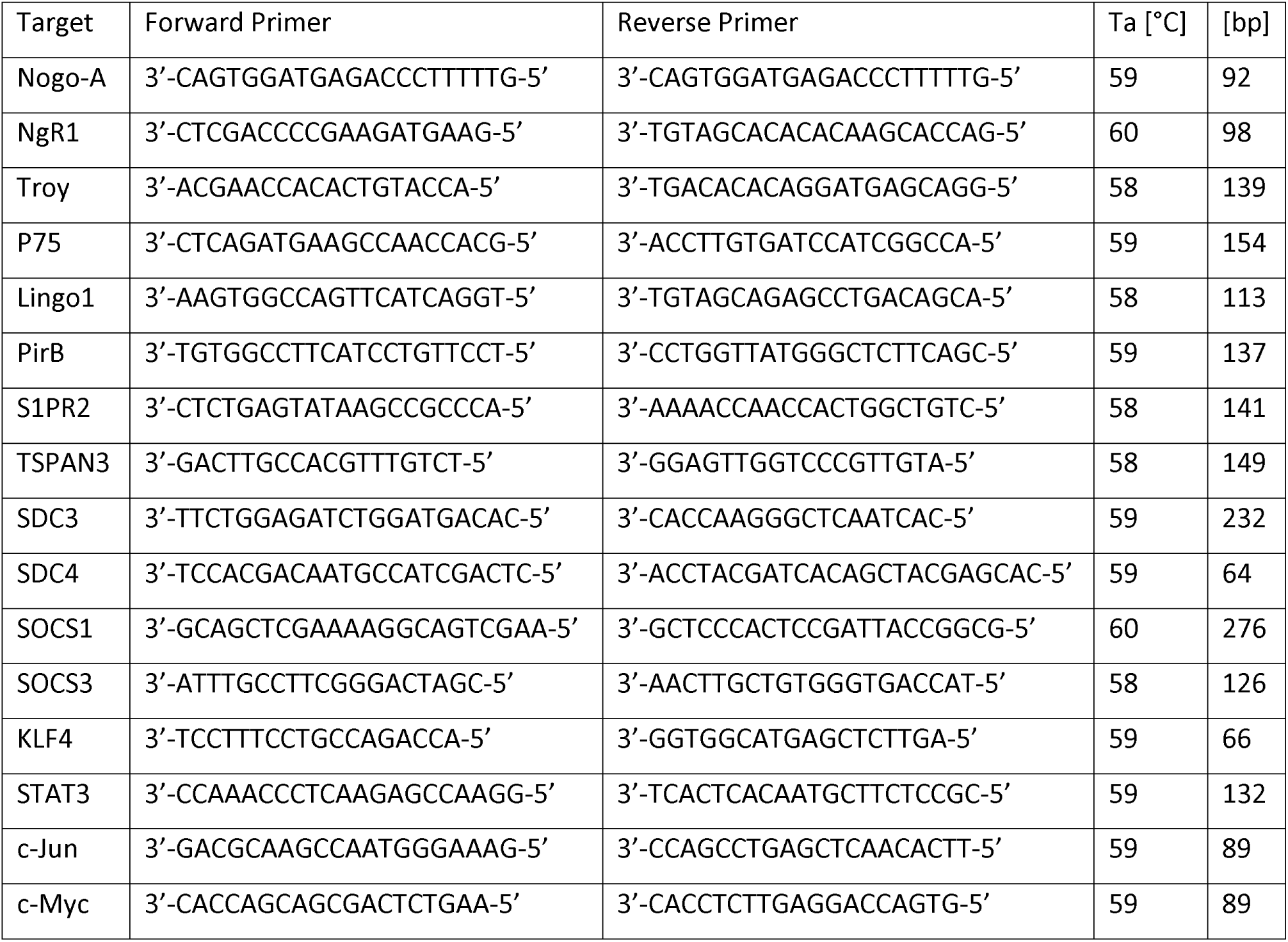
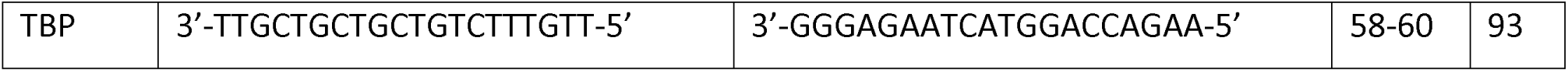
Primer sequences used for qRT-PCR.

### Immunocytochemistry

Fixed cells were blocked and permeabilized in blocking buffer (0.1% Triton-X, 10% BSA in PBS) for 1 h. Primary antibodies were applied in 1% BSA in PBS and incubated over night at 4°C. The samples were washed three times in PBS, followed by incubation with secondary antibodies in 1% BSA in PBS for 1 h at RT. After washing three times in PBS, the samples were coverslipped in fluorescence mounting medium (Dako) and imaged on an Axioskop2 fluorescence microscope (Zeiss) equipped with a Plan NEOFLUAR dry 20x/0.5 objective and Zeiss Axiocam MRc camera using the Zeiss Axiovision software. For image acquisition, exposure times were kept constant and below grey scale saturation. For immunofluorescence measurement, the signal in the phalloidin channel was thresholded and an area mask was created around the fluorescent object using the ImageJ software (NIH, Bethesda, MD, USA). This mask was then applied onto the RhoA or p-ERK1/2 pictures and the total pixel intensity within the area was measured. This value was normalized to the area of the growth cone or cell body. Similarly, the nuclear counter stain was used to define the area of the nucleus. Each experiment was conducted in three technical replicates and all quantifications show at least three independent experiments. Per technical replicate fife randomly chosen images were analyzed, which contained between 10 and 20 cells.

### Statistics

All graphs and statistical analyses were computed in GraphPad Prism version 7.03 (GraphPad Software Inc., La Jolla, CA, USA). All data were normalized to baseline values and plotted as mean values ± standard error of the mean (SEM). One-way analysis of variance (ANOVA) with Bonferroni multiple comparisons test was used to compare multiple groups and Dunnet multiple comparison test was used to compare multiple groups to the baseline (only in the analysis of Nogo-A and receptor expression during differentiation of N1E-115 cells). Asterisks indicate the following p-values: *p < 0.05, **p < 0.01, ***p < 0.001.

## Results

### N1E-115 Neuroblastoma Cells as a Model System to Study neurite outgrowth and Nogo-A Signaling

We characterized the expression of Nogo-A and its receptors in the mouse neuroblastoma-derived cell line N1E-115. In the absence of serum, this cell line readily differentiates into a neuron-like morphology within 24 hours and continues to form processes thereafter (Sup. Fig.1 A, B). In the undifferentiated state, as well as in the 24 and 48 hour differentiated states, the cells expressed all known Nogo-A receptors (Kempf et al., 2017; Thiede-Stan et al., 2015; Kempf et al., 2014; Kempf and Schwab, 2013)) (Sup. Fig.1 C, D). To model the growth inhibitory CNS environment *in vitro*, we treated the N1E-115 cells with crude spinal cord extract (SCE) containing soluble as well as membrane proteins including myelin proteins. Three independent preparations of SCE contained a comparable level of Nogo-A (1.33 ± 0.05 nM) (Sup. Fig.2 A-C) and treatment of 24 h differentiated cells with SCE inhibited subsequent neurite outgrowth in a dose dependent manner (Sup. Fig.2 D, E).

**Fig. 1.**
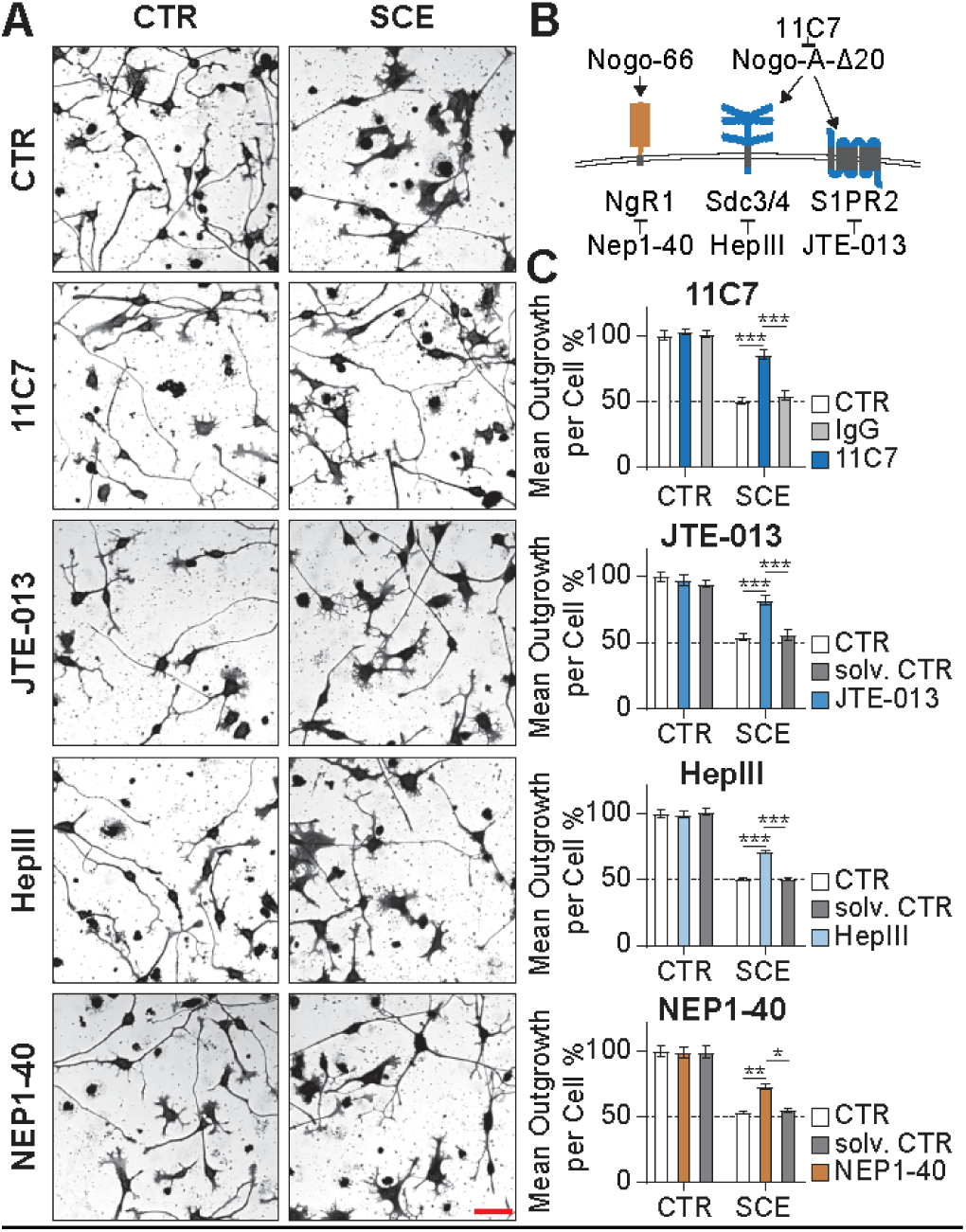
Inhibition of key Nogo-A signaling components rescues N1E-115 neurite outgrowth in presence of growth inhibitory spinal cord extract. (A) Representative pictures of N1E-115 outgrowth inhibition by spinal cord extract (SCE) and rescue thereof through treatment with function-blocking Nogo-A antibody (11C7), S1PR2 antagonist (JTE-013), heparinase III (HepIII) and NgR1 blocking peptide (NEP1-40). (B) Schematic representation of the examined inhibitors of Nogo-A signaling components. (C) Quantification of mean N1E-115 outgrowth per cell in the different treatment conditions. Red scale bar = 50 µm, n = 5 independent experiments, *p ≤ 0.05, **p ≤ 0.01, ***p ≤ 0.001.

Next, we investigated whether part of the SCE-mediated neurite outgrowth inhibition can be attributed to Nogo-A signaling by blocking the Nogo-A-Δ20 signaling domain as well as several Nogo-A receptors (Fig.1 B). Direct inhibition of Nogo-A-Δ20 via the function-blocking antibody 11C7 resulted in a robust rescue of neurite outgrowth in presence of SCE, compared to the control antibody condition (Fig.1 A, C). Similarly, co-treatment with SCE and JTE-013, a functional antagonist of the Nogo-A-Δ20 receptor S1PR2 (Marsolais and Rosen, 2009), and enzymatic cleavage of heparin sulfates from the cell surface to inhibit the Nogo-A-Δ20 receptors Sdc3/4 (Kempf et al., 2017) also increased neurite outgrowth when compared to the SCE and solvent control condition (Fig.1 A, C). In addition, blockade of the Nogo-66 receptor NgR1 via the antagonist peptide NEP1-40 (GrandPré et al., 2002) alleviated the SCE-mediated inhibition of neurite outgrowth (Fig.1 A, C). These results demonstrate that a substantial part of the SCE-mediated outgrowth inhibition is mediated by Nogo-A. Interestingly, the Nogo-A-Δ20 interacting receptors S1PR2 and Sdc3/4 as well as the Nogo-66 binding NgR1 are involved in this growth restricting signaling event.

### Nogo-A Activates ERK1/2 and mTORC1 Downstream Effector S6K

Multiple studies have reported mTORC1 as a downstream target of Nogo-A (Manns et al., 2014; Peng et al., 2011; Raiker et al., 2010; Gao et al., 2010; Wang et al., 2008). Therefore, we tested whether SCE differentially regulated either the mTORC1 activating upstream protein AKT, or the downstream proteins elongation and initiation factor 4E binding protein 1 (4E-BP1) and ribosomal protein S6 kinase (S6K) (Saxton and Sabatini, 2017). SCE treatment induced AKT phosphorylation on S473 suggesting differential mTORC2 activation (Fig.2 A, B). However, many neurotrophic factors induce AKT phosphorylation at T308, and this phosphorylation site has been associated with the pro-regenerative effects of AKT overexpression *in vivo* (Keefe et al., 2017; Miao et al., 2016). Furthermore, possible signaling crosstalk via Nogo-A downstream effector ROCK mediated activation of PTEN, which results in negative regulation of AKT at the T308 site, was suggested in the context of cellular chemotaxis (Li et al., 2005). While SCE treatment did not result in decreased phospho-T308 AKT levels (Fig.2 A, B), SCE co-treatment with insulin decreased the insulin mediated upregulation of phospho-T308 AKT (Fig.2 C, D). These results suggest that depending on cellular context, SCE affects AKT differentially on its two phosphorylation sites.

Next, we investigated whether this differential AKT phosphorylation propagated to the activation of mTORC1 downstream proteins. While phosphorylation of 4E-BP1 at S65 or T37/46 was not altered by SCE treatment, SCE induced a drastic increase in S6K and the downstream ribosomal protein S6 (S6P) phosphorylation (Fig.2 E, F). This contradictory result suggests a lateral mTORC1 activating signal integration downstream of AKT, or alternatively a direct S6K activation by factors in the SCE, e.g. via ERK1/2 or RSK.

**Fig. 2.**
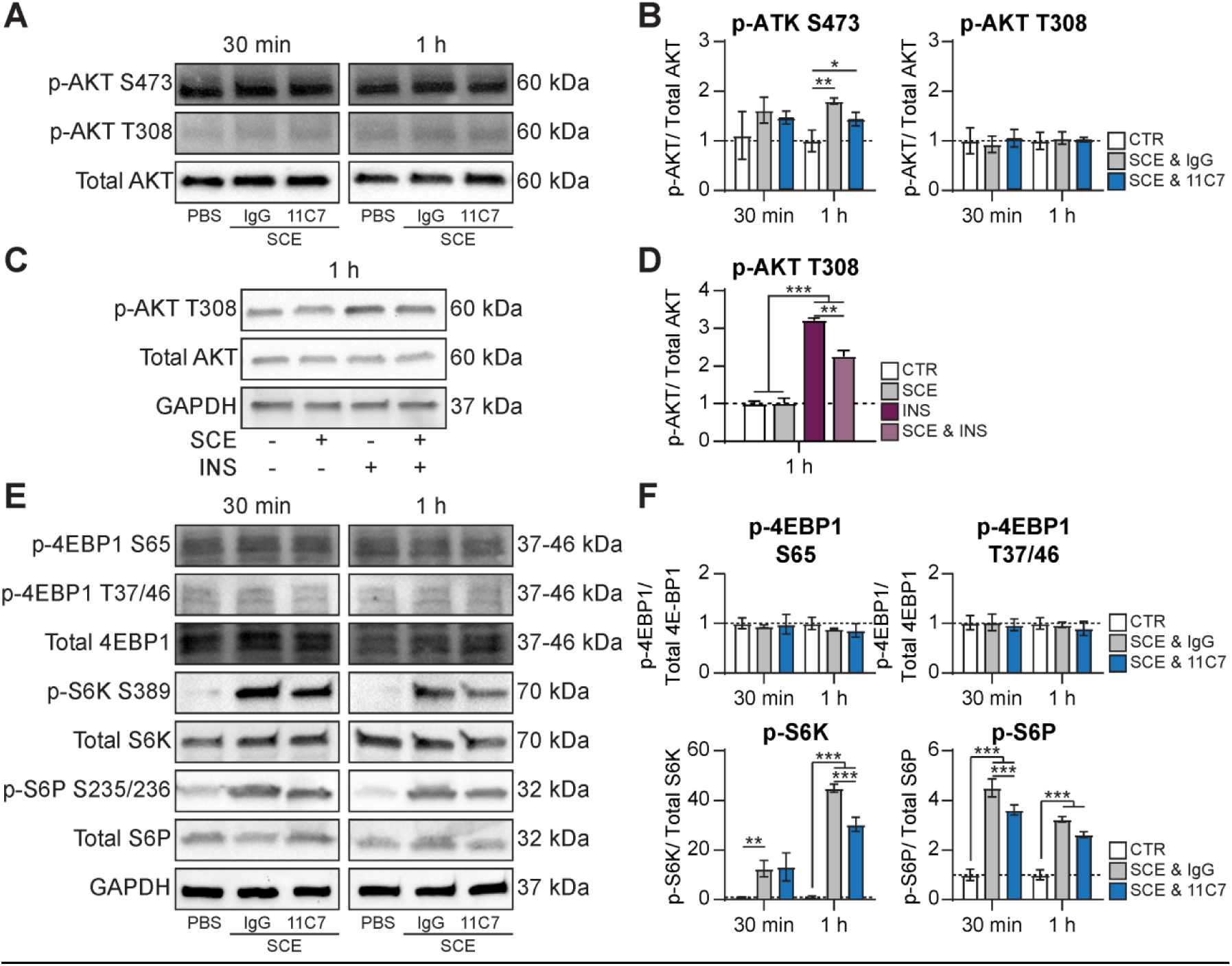
SCE downstream signaling integrates into the AKT/mTORC1 pathway. (A, B) Representative p-AKT S473, p-AKT T308, and AKT western blots of N1E-115 cells treated with SCE in combination with either control antibody (IgG) or Nogo-A-Δ20 function-blocking antibody (11C7) and quantification thereof. (C, D) Representative western blots of cells treated with either insulin and SCE alone or in combination and quantification thereof. (E, F) Representative p-4E-BP1 S65, p-4E-BP1 T37/46, 4E-BP1, p-S6K S389, S6K, p-S6P S235/236 and S6P western blots of N1E-115 cells treated with SCE in combination with either control antibody (IgG) or Nogo-A-Δ20 function-blocking antibody (11C7) and quantification thereof. n = 3 independent experiments, *p ≤ 0.05, **p ≤ 0.01, ***p ≤ 0.001.

The mitogen activated protein kinase (MAPK) ERK1/2 can activate mTORC1 downstream of AKT via inhibitory phosphorylation of tuberous sclerosis complex 1/2 (TSC1/2). Furthermore, it can lead to activation of S6K via direct phosphorylation by the downstream kinase ribosomal S6 kinase (RSK) (Saxton and Sabatini, 2017). Treatment of N1E-115 cells with SCE resulted in pronounced ERK1/2 activation within 30 min to 1 h after treatment, as reflected by phosphorylation of the sites T202/Y204 in its activation loop, as well as phosphorylation of its downstream effector RSK (Fig.3 A-D). This activation was partially Nogo-A dependent, as inhibition of the Nogo-A-Δ20 domain via function-blocking antibodies resulted in a decrease of ERK1/2 phosphorylation. Furthermore, pharmacological inhibition of ERK1/2 with the small molecule SCH772984 (Morris et al., 2013) partially rescued the SCE-mediated decrease in neurite outgrowth, suggesting that ERK1/2 activation is a downstream effect of the inhibitory ligands in the SCE (Fig.3 F, G).

**Fig. 3.**
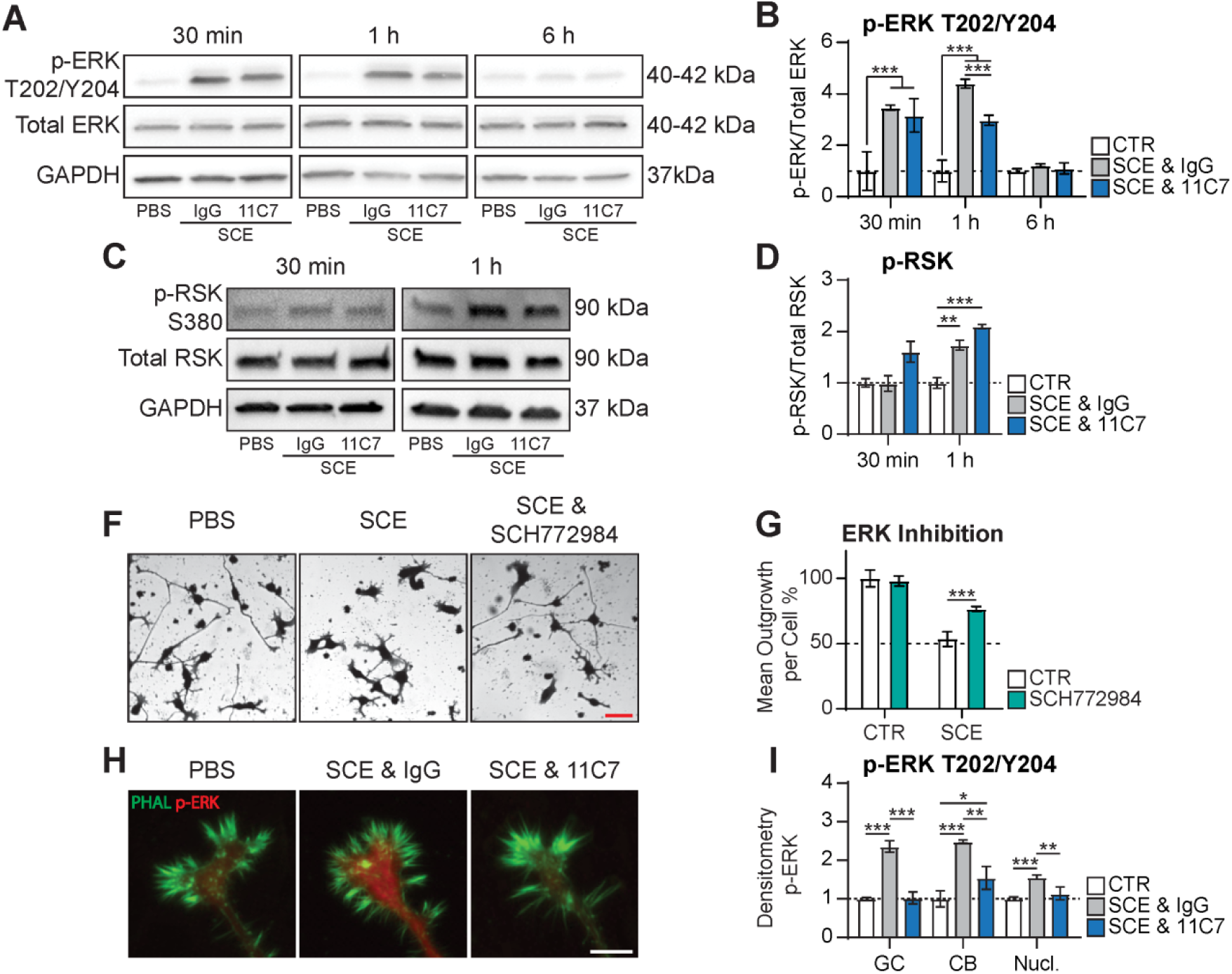
Inhibitory effect of SCE is partially dependent on ERK1/2 activation. (A, B) Representative western blots for p-ERK1/2 and total ERK1/2 of N1E-115 cells treated with SCE in combination with either control antibody (IgG) or Nogo-A-Δ20 function-blocking antibody (11C7) and quantification of activated ERK1/2 (p-ERK1/2/total ERK1/2). (C, D) Representative western blots for p-RSK and total RSK of N1E-115 cells treated with SCE in combination with either control antibody (IgG) or Nogo-A-Δ20 function-blocking antibody (11C7) and quantification thereof. (F, G) Representative pictures of N1E-115 cells treated for 24 h with SCE only or SCE in combination with ERK inhibitor (SCH772984) and quantification of mean outgrowth per cell thereof. (H) Representative pictures of growth cones stained for p-ERK1/2 (red) and f-actin (phalloidin; green) of N1E-115 cells treated for 1 h with SCE in combination with either control antibody (IgG) or Nogo-A-Δ20 function-blocking antibody (11C7). (I) Quantification of activated ERK1/2 in the growth cone (GC), the cell body (CB) and the nucleus (Nucl.). Red scale bar = 50 µm, white scale bar = 5 µm, n = 3 independent experiments for WB, n = 5 independent experiments for outgrowth assays, n = 3 independent experiments for IF, *p ≤ 0.05, **p ≤ 0.01, ***p ≤ 0.001.

ERK1/2 mediated signaling is associated with diverse functions, and its localization is regulated by a plethora of adaptor and scaffolding proteins to ensure a precise spatiotemporal signaling response (Plotnikov et al., 2010). Therefore, we investigated the cellular location of the active phospho-ERK1/2. While ERK1/2 was activated by SCE globally in our model system, a more pronounced activation was observed in the growth cones and the cell body, suggesting a role apart from transcriptional control (Fig.3 H, I). Importantly, the activation of ERK1/2 was completely blocked by 11C7 treatment in the growth cone, further corroborating ERK1/2 activation as a downstream target of Nogo-A. Taken together, these results identify ERK1/2 as a novel Nogo-A signaling effector in neurite outgrowth inhibition and growth cone collapse.

### Nogo-A Induces Local Protein Synthesis of RhoA via an ERK1/2 Dependent Pathway

Several studies have demonstrated that the regenerative capacity of some axons depends on local protein synthesis and degradation in the growth cone (Gumy et al., 2010). Intriguingly, also growth cone collapse mediated by inhibitory cues, such as semaphorins and Nogo-A, were shown to be dependent on local protein synthesis (Manns et al., 2014; Wu et al., 2005). Both protein translation and degradation are processes that have been associated with MAPK signaling (Roux and Topisirovic, 2012). Based on the increase in phospho-S6K and phospho-S6P and the observed Nogo-A dependent ERK activation in the growth cone, we hypothesized that local protein synthesis of Nogo-A downstream proteins could enhance Nogo-A mediated axonal growth inhibition. Therefore, we analyzed the total levels of RhoA, an integrator of many growth inhibitory cues including Nogo-A (Kempf and Schwab, 2013). Total RhoA levels increased in N1E-115 cells after treatment with SCE; this effect was prevented by the addition of Nogo-A neutralizing antibodies (Fig.4 A, B). Next, we tested whether the activation of S6P and RhoA translation are downstream effects of ERK1/2 signaling. While activation of S6P was partially reduced by ERK1/2 inhibition, the RhoA increase was completely abolished by the specific ERK1/2 inhibitor SCH772984 (Fig.4 C-G). These results show that in addition to activating the RhoA/ROCK enzymatic cascade, Nogo-A can further reduce neuronal growth and regeneration by upregulating RhoA levels in growth cones via an ERK1/2 dependent pathway.

**Fig. 4.**
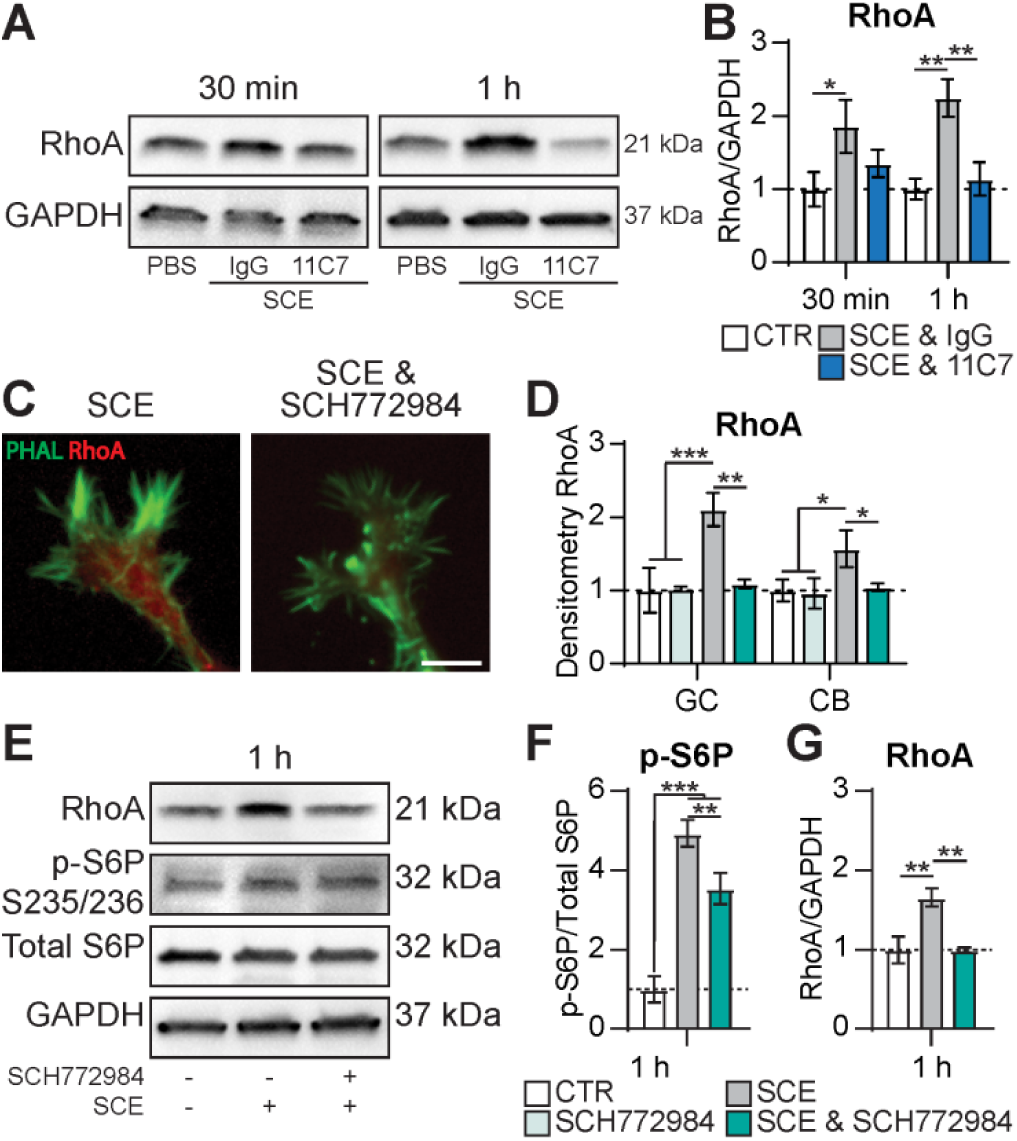
Nogo-A elevates RhoA protein levels via an ERK1/2 dependent pathway. (A, B) Representative RhoA western blot of N1E-115 cells treated with SCE in combination with either control antibody (IgG) or Nogo-A-Δ20 function-blocking antibody (11C7) and quantification thereof. (C) Representative pictures of growth cones stained for RhoA (immunofluorescence (IF); red) and f-actin (phalloidin; green) of N1E-115 cells treated for 1h with SCE in combination with either solvent control or ERK1/2 inhibitor (SCH772984). (D) Quantification of RhoA in growth cones (GC) and cell bodies (CB). (E-G) Representative RhoA, p-S6P and total S6P western blots of N1E-115 cells treated with SCE in combination with either solvent control or ERK1/2 inhibitor (SCH772984) and quantification thereof. White scale bar = 5 µm, n = 3 independent experiments for WB, n = 3 independent experiments for IF, *p ≤ 0.05, **p ≤ 0.01, ***p ≤ 0.001.

### Nogo-A Upregulates JAK/STAT Pathway Inhibitors SOCS3 and KLF4

Apart from the growth cone, ERK1/2 activation was also induced by Nogo-A or SCE in the neuronal soma and to a lesser extent in the nucleus. ERK1/2 has been described to alter the transcriptional profile in the nucleus (Plotnikov et al., 2010). The transcription factor STAT3 has been previously shown to be upregulated in Nogo-A antibody treated rat brains (Bareyre et al., 2011), to mediate the conditioning lesion effect in DRG neurons (Qiu et al., 2005) and to boost axonal regeneration following optic nerve crush (Mehta et al., 2016; Pernet et al., 2013). STAT proteins are regulated by two distinct phosphorylation events. Tyrosine phosphorylation in the SH2 domain by JAK induces dimerization and nuclear translocation, a process which can be modulated by ERK1/2-mediated serine phosphorylation in the STAT linker domain (Tian and An, 2004; Jain et al., 1998; Wen et al., 1995). Phospho-profiling of SCE treated N1E-115 cells revealed a rapid increase in phosphorylation at the ERK1/2 associated phospho-site (Fig.5 A, B), while no changes could be detected at the JAK2-associated site (data not shown). Previous work has indicated that exclusive phosphorylation of the STAT linker domain negatively regulates tyrosine phosphorylation and STAT mediated transcription (Tian and An, 2004; Jain et al., 1998). In fact, SCE-treatment induced a depletion of total STAT3 levels, suggesting a regulatory mechanism for direct negative effects of neurite growth inhibitory factors on the JAK/STAT pathway (Fig.5 C). In contrast, activation of tyrosine STAT3 phosphorylation induced by interleukin 6 (IL-6) treatment (Schumann et al., 1999) rescued neurite outgrowth in the SCE treated condition (Fig.5 D,E), making STAT3 a potential integration point for neurite outgrowth-promoting and outgrowth-inhibiting signals.

**Fig. 5.**
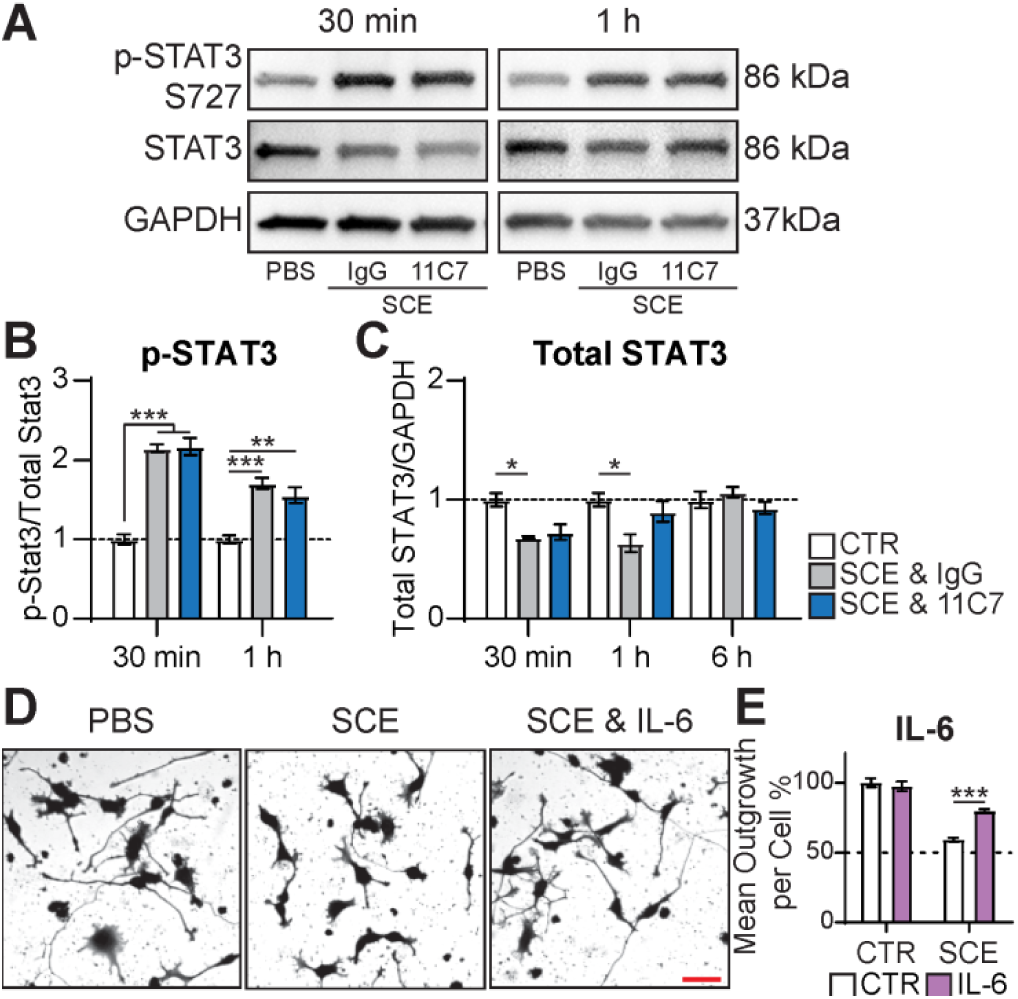
STAT3 as an integration point for neurite outgrowth inhibitory and promoting signals. (A-C) Representative p-STAT3 and total STAT3 western blots of N1E-115 cells treated with SCE in combination with either control antibody (IgG) or Nogo-A-Δ20 function-blocking antibody (11C7) and quantification thereof. (D, E) Representative pictures of N1E-115 cells treated with SCE alone or SCE in combination with IL-6 and quantification of mean outgrowth per cell thereof. Red scale bar = 50 µm, n = 5 independent experiments for outgrowth assays, n = 3 independent experiments for WB, *p ≤ 0.05, **p ≤ 0.01, ***p ≤ 0.001.

We further tested whether Nogo-A interferes with the JAK/STAT pathway by regulating its well characterized inhibitors SOCS1, SOCS3 and KLF4. We observed a Nogo-A dependent upregulation of SOCS3 mRNA levels after 6 h and SOCS1 and KLF4 mRNA levels after 24 h (Fig.6 A-C). The observed early Nogo-A independent decrease in SOCS3 and KLF4 mRNA levels could correspond to the decrease of STAT3 protein levels described above, as they are transcriptional targets of the JAK/STAT pathway themselves. Furthermore, both SOCS3 and KLF4 protein levels were upregulated in a Nogo-A dependent manner upon SCE treatment of N1E-115 cells (Fig.6 D-F). Interestingly, SOCS3 protein levels increased before it was transcriptionally upregulated, suggesting a dual mechanism in its regulation. These results demonstrate a novel signaling route via STAT pathway inhibition by which Nogo-A could decrease axonal growth and regeneration.

**Fig. 6.**
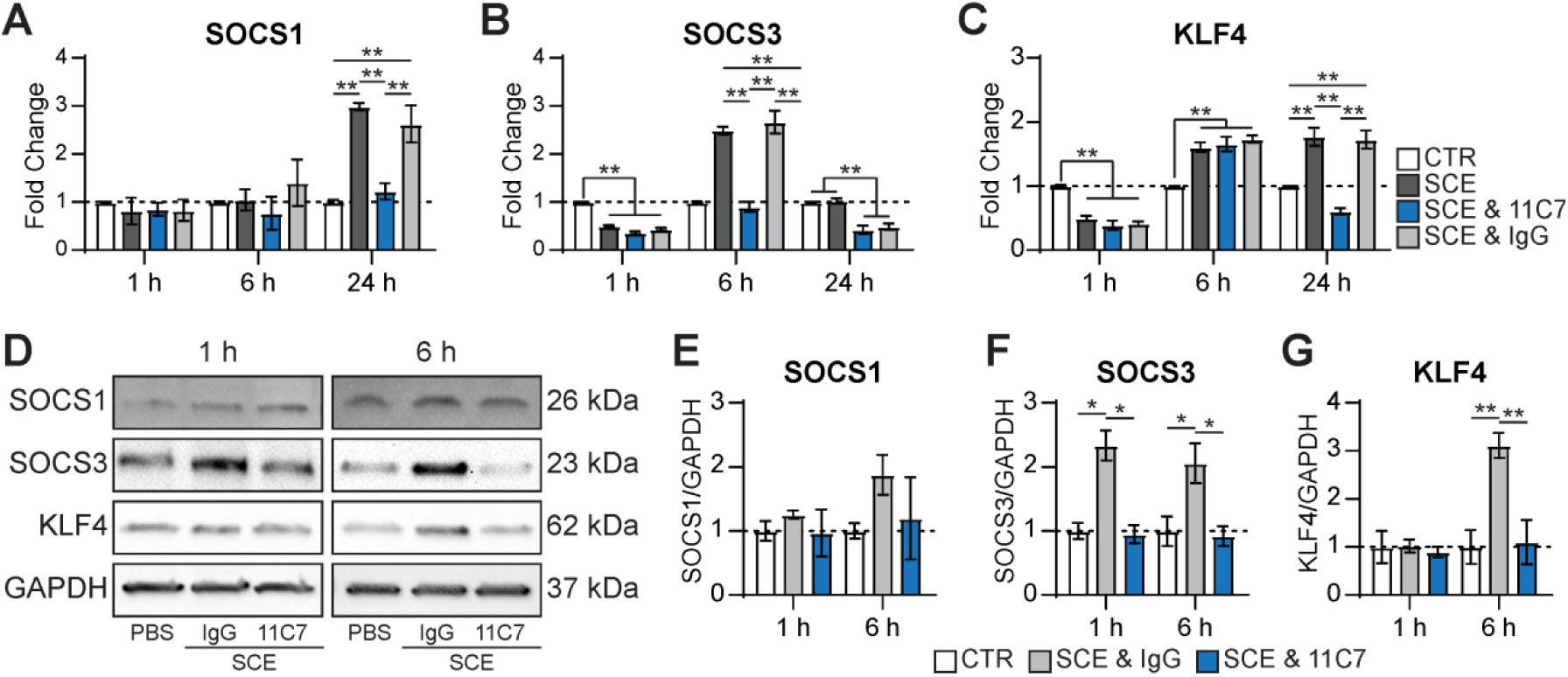
Nogo-A elevates levels of JAK/STAT pathway inhibitors SOCS3 and KLF4. (A-C) Transcriptional changes of SOCS1, SOCS3 and KLF4 in N1E-115 cells treated with either SCE alone or SCE in combination with control antibody (IgG) or Nogo-A-Δ20 function-blocking antibody (11C7). (D - G) Representative SOCS1, SOCS3 and KLF4 western blots of N1E-115 cells treated with SCE in combination with either control antibody (IgG) or Nogo-A-Δ20 function-blocking antibody (11C7) and quantification thereof. n = 3 independent experiments for qRT-PCR and WB, *p ≤ 0.05, **p ≤ 0.01, ***p ≤ 0.001.

## Discussion

Using a robust neurite outgrowth assay with N1E-115 mouse neuroblastoma cells, we show that spinal cord extracts induce a differential activation of the mTORC1 effectors S6K and 4E-BP1 downstream of AKT, likely via activation of the MAPK ERK1/2 in a Nogo-A dependent way. Rapid activation of the ERK1/2 pathway by Nogo-A and SCE mediates the local synthesis and accumulation of RhoA in growth cones thereby potentiating Nogo-A mediated growth cone collapse. Furthermore, Nogo-A signaling could directly inhibit the JAK/STAT pathway via upregulation of its inhibitors SOCS3 and KLF4.

Over the last two decades, several Nogo-A receptors have been identified, showing complex multi-ligand multi-receptor signaling system (Kempf et al., 2017; Thiede-Stan et al., 2015; Kempf et al., 2014; Kempf and Schwab, 2013). This is the first study utilizing a cell line that shows expression of all Nogo-A receptors, thereby corroborating the importance of several key receptors in one functional assay. In the past, many *in vitro* studies have focused on a single ligand receptor interaction in the context of axonal growth or the lack thereof. However, a series of studies highlight the importance of investigating cellular signaling as an integrative, context-dependent process (Mazel, 2017; Day et al., 2016). Our approach of treating cells with crude spinal cord extract (SCE) instead of selected ligands offers the advantage of modelling the *in vivo* situation more closely. By applying specific blockers, in particular a monospecific antibody against Nogo-A, along with the SCE, we were able to delineate the individual contributions of specific inhibitory cues within the SCE. Furthermore, the use of a cell line over primary neuronal cultures in our neurite outgrowth assay allows for both a faster and more cost effective screening and exploration of potential novel outgrowth promoting interventions in an environment that resembles the physiological situation within the CNS.

Previous studies have indicated crosstalk between Nogo-A signaling and the AKT/mTORC1 pathway, while the underlying mechanisms remained unclear (Gao et al., 2010; Manns et al., 2014; Peng et al., 2011; Raiker et al., 2010; Sun et al., 2015; Wang et al., 2008). *In vitro* evidence for Nogo-A downstream effector ROCK dependent phosphorylation and thereby activation of PTEN could represent one possible signaling internode for such a crosstalk (Li et al., 2005). Consistently, SCE treatment decreased insulin-induced phosphorylation on the T308 site of AKT in our study. Interestingly, the mTORC2-associated S473 site of AKT was upregulated after treatment with SCE suggesting an activation of mTORC2 by factors within the SCE. Consensus in the literature is that full activation of AKT requires both phosphorylation events (Saxton and Sabatini, 2017), although a recent study by (Miao et al., 2016) reported that the two phosphorylation sites T308 and S473 affect neurite regeneration in opposing ways: While the canonical PDK1/AKT/mTORC1 pathway had a positive impact on regeneration, phosphorylation of AKT on S473 blocked regeneration, possibly by activating the mTORC1 inhibitor TSC1/2 via GSK3.

Phosphorylation of the mTORC1 effector S6K and downstream S6P was induced by SCE treatment, partially in a Nogo-A dependent way. This indicates lateral integration of Nogo-A signaling in the mTORC1 pathway downstream of AKT, possibly via activation of the MAPK ERK1/2 and downstream effector RSK, which were highly activated by SCE treatment. The MEK/ERK1/2 pathway and AKT/mTORC1 pathways have been associated with substantial signaling crosstalk on multiple levels. While MEK dependent phosphorylation of AKT inhibits the upstream pathway, ERK1/2 or RSK dependent phosphorylation of TSC1/2 or mTORC1 adaptor protein raptor activates mTORC1 signaling (Saxton and Sabatini, 2017). Additionally, this downstream integration of inhibitory signaling might in turn further inhibit the AKT upstream PI3K by activating phosphorylation-dependent degradation of its scaffolding subunit IRS1 (Saxton and Sabatini, 2017), thereby desensitizing the cell for further growth promoting extrinsic cues.

SCE treatment induced rapid Nogo-A dependent activation of ERK1/2 in the growth cones, and global inhibition of ERK1/2 activation rescued neurite outgrowth upon SCE treatment. Furthermore, we observed an ERK1/2-dependent increase in RhoA levels upon SCE-treatment similar to a previously published study in the context of semaphorin 3A mediated growth cone collapse (Wu et al., 2005). Several studies have linked ERK1/2 activation and retrograde transport with induction of pro-regenerative transcriptional changes downstream of neurotrophic factor signaling (O’Donovan et al., 2014; Perlson et al., 2006; Chao, 2003) suggesting a dual role for ERK1/2 signaling dependent on its subcellular localization. A similar location-dependent role in axonal regeneration was described for DLK1 in the context of retinal ganglion cell axon regeneration: Following optic nerve crush, DLK1 was responsible for the majority of injury-elicited gene expression changes, and its deletion enhanced retinal ganglion cell survival (Watkins et al., 2013). However, co-deletion of DLK1 and PTEN ablated the strong pro-regenerative effect of PTEN deletion after optic nerve crush, suggesting a positive role for DLK1 in the induction of regenerative sprouting. Possible differential regulation of ERK1/2 scaffolding proteins, the machinery mediating retrograde transport, and the underlying mechanism of activation remain to be investigated. Furthermore, the internalization and retrograde transport of recombinant Nogo-A-Δ20 fragments was reported in multiple studies (Thiede-Stan et al., 2015; Kempf et al., 2014; Joset et al., 2010), however, it remains to be investigated whether this also occurs with endogenous full length Nogo-A, and how the signaling effectors of internalized and cell surface Nogo-A might differ.

SCE treatment of the neuronal cells resulted in negative JAK/STAT pathway regulation on several levels. We observed increased, potentially ERK1/2 mediated serine phosphorylation of STAT3 as well as STAT3 depletion. Furthermore, SCE treatment resulted in a Nogo-A dependent upregulation of the STAT3-inhibitors SOCS3 and KLF4 both on a transcriptional and translational level. An earlier study (Miao et al., 2006) identified SOCS3 as a negative regulator of STAT3 tyrosine phosphorylation and nuclear translocation, resulting in reduced neurite outgrowth. Later studies found that SOCS3 deletion in retinal ganglion cells induced robust axonal regeneration following optic nerve crush (Smith et al., 2009), while SOCS3 overexpression reduced optic nerve regeneration (Hellström et al., 2011). It is therefore intriguing that Nogo-A induced SOCS3 expression and may thereby inhibit neuronal outgrowth via direct inhibition of the JAK/STAT cascade. Also KLF4 has been shown to repress axonal growth both *in vitro* and *in vivo* via inhibition of STAT3 (Qin et al., 2013; Moore et al., 2009). In the optic nerve, a marked increase in KLF4 expression was reported around birth (Moore et al., 2009), a time point also associated with the invasion of oligodendrocyte precursor cells into the optic nerve (Pernet et al., 2008). However, it remains to be investigated whether there is a functional link between KLF4 expression in neuronal cells and Nogo-A expression in oligodendroglial cells.

The results presented in this study provide novel insights into Nogo-A mediated intracellular signaling beyond the activation of the RhoA/ROCK pathway. We show that the MAPK ERK1/2 is activated by Nogo-A and may directly affect growth cone dynamics via regulation of local protein synthesis, in particular of RhoA. Furthermore, we highlight several novel transcriptional regulators which were affected by Nogo-A signaling on a transcriptional, translational as well as a post-translational level. Finally, we demonstrate a possible inhibitory crosstalk between Nogo-A signaling and the growth promoting AKT/mTORC1 and JAK/STAT pathways. These data highlight the importance of studying signaling as an integrative process and further promote our understanding of how neurite outgrowth inhibitory cues are integrated into the neuronal signaling network.

## Acknowledgements

This work was supported by the Swiss National Science Foundation (grant Nr. 31-138676/1) and an advanced ERC grant (NOGORISE) to the senior author.

## Author contributions

M. Maibach designed the research and wrote the manuscript and designed the figures. M. Maibach, E. Piovesana and contributed to data acquisition and analysis. E. Piovesana, J. Kaiser, M. Maurer and M.E. Schwab gave valuable input in experimental and manuscript discussions.

## Conflict of interest

The authors declare no competing financial interests.

## Figure legends

**SupFig. S1.**
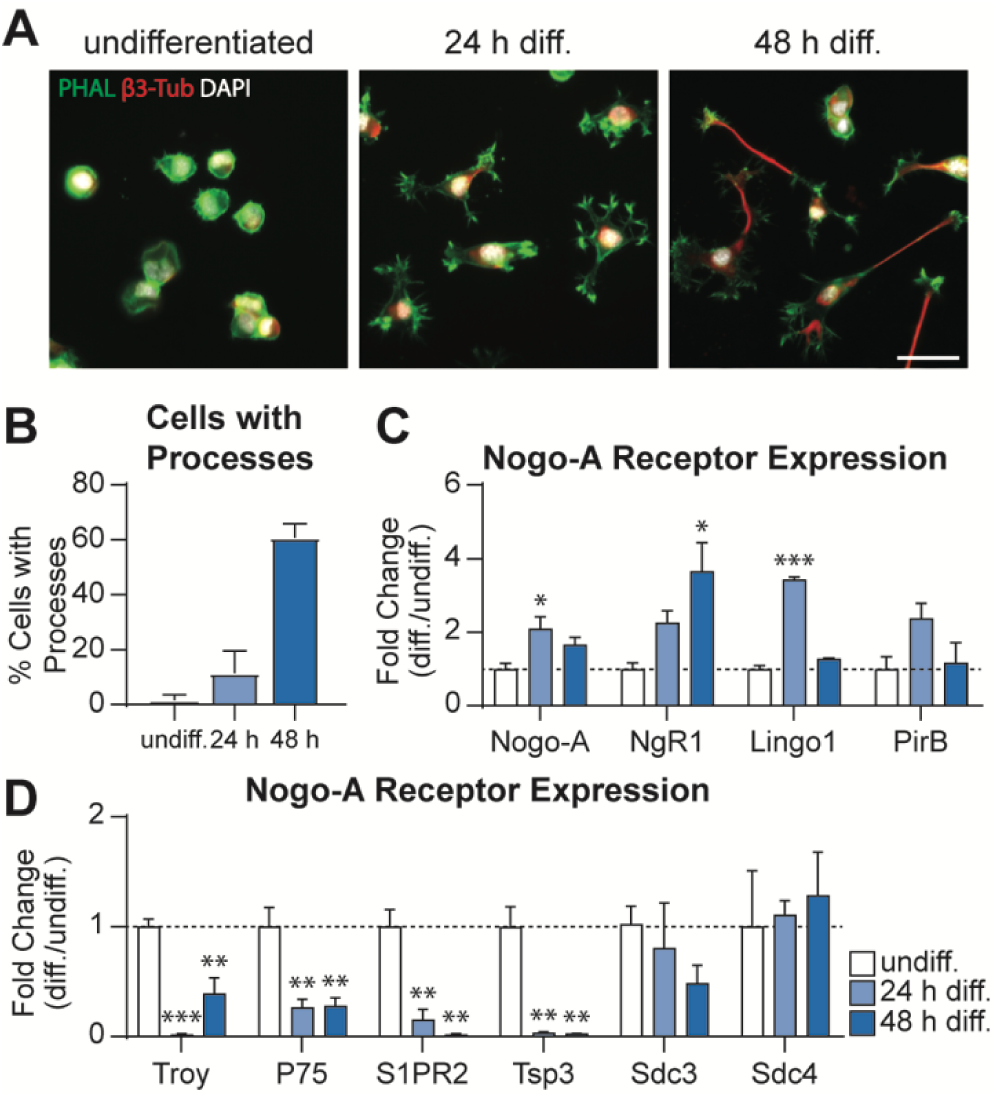
N1E-115 cells as a model system to study Nogo-A mediated neurite outgrowth inhibition. (A) Representative pictures of undifferentiated, 24 h and 48 h differentiated N1E-115 cells stained for f-actin (phalloidin; green) and β3-tubulin (red). (B) Quantification of cells with processes longer than the cell diameter. (C, D) Transcriptional analysis of Nogo-A and Nogo-A receptors in undifferentiated, 24 h and 48 h differentiated N1E-115 cells. Asterisks indicate significant changes relative to the undifferentiated group. White scale bar = 50 µm, n = 3 independent experiments, *p = 0.05, **p = 0.01, ***p = 0.001.

**SupFig. S2.**
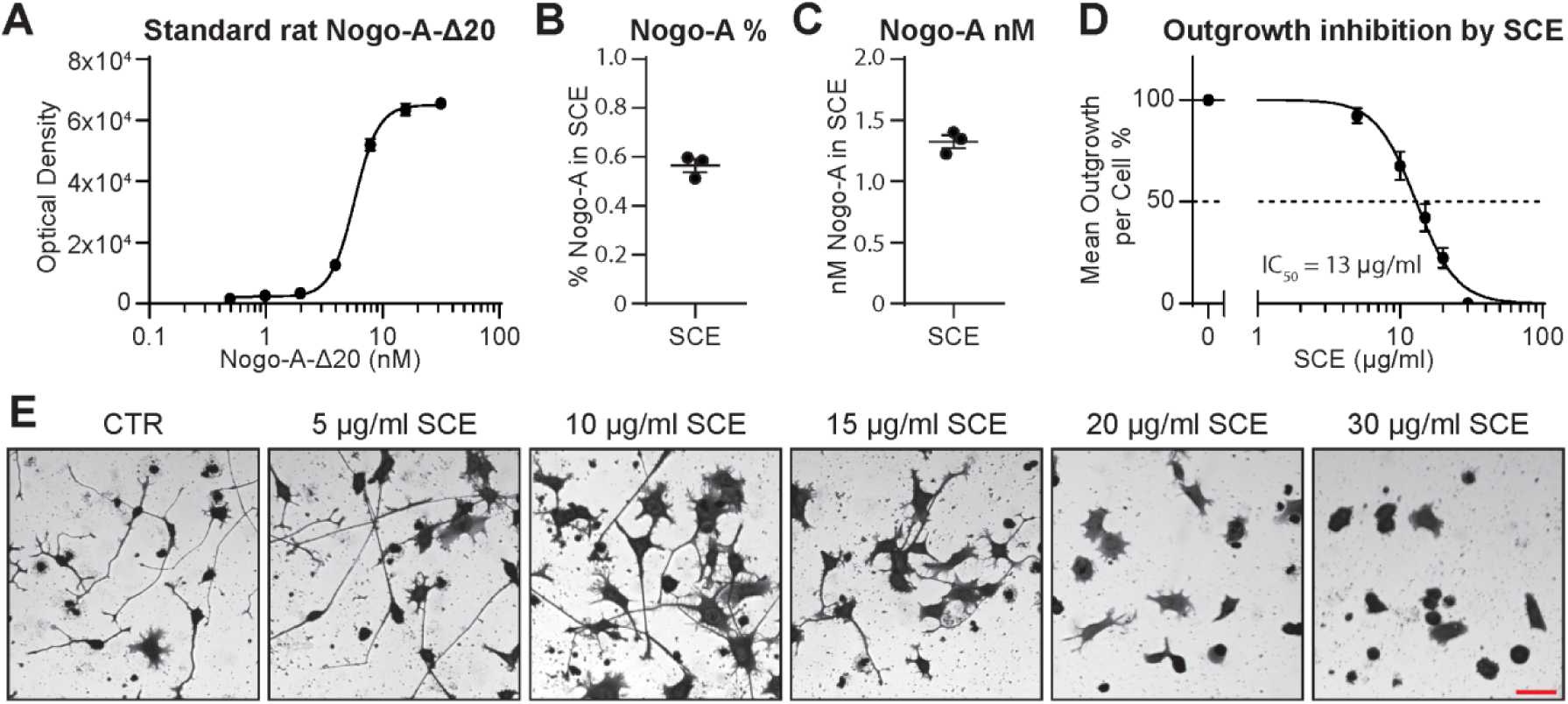
Spinal cord extract contains Nogo-A and inhibits neurite outgrowth in a dose-dependent manner. (A-C) Representative standard curve of Nogo-A-Δ20 (A) used to calculate the percentage of Nogo-A (B) and nM concentration of Nogo-A (C) in three independent spinal cord extract (SCE) preparations. (D) SCE inhibits outgrowth of N1E-115 cells in a dose dependent manner. The mean IC_50_ of three independent SCE preparations was 13 µg/ml. (E) Representative pictures of N1E-115 neurite outgrowth inhibition by increasing SCE concentrations. Red scale bar = 50 µm, n = 3 independent experiments.

## Notes

### Competing Interest Statement

The authors have declared no competing interest.

